# Fatecode: Cell fate regulator prediction using classification autoencoder perturbation

**DOI:** 10.1101/2022.12.16.520772

**Authors:** Mehrshad Sadria, Anita Layton, Sidharta Goyal, Gary D. Bader

## Abstract

Cell reprogramming, which guides the conversion between cell states, is a promising technology for tissue repair and regeneration. Typically, a group of key genes, or master regulators, are manipulated to control cell fate, with the ultimate goal of accelerating recovery from diseases or injuries. Of importance is the ability to correctly identify the master regulators from single-cell transcriptomics datasets. To accomplish that goal, we propose Fatecode, a computational method that combines in silico perturbation experiments with cell trajectory modeling using deep learning to predict master regulators and key pathways controlling cell fate. Fatecode uses only scRNA-seq data from wild-type samples to learn and predict how cell type distribution changes following a perturbation. We assessed Fatecode’s performance using simulations from a mechanistic gene regulatory network model and diverse gene expression profiles covering blood and brain development. Our results suggest that Fatecode can detect known master regulators of cell fate from single-cell transcriptomics datasets. That capability points to Fatecode’s potential in accelerating the discovery of cell fate regulators that can be used to engineer and grow cells for therapeutic use in regenerative medicine applications.

## Introduction

In tissue development, specific regulator genes play a key role in determining how cells decide to change state and type to form a complete tissue with many different cell types. These regulators are also important in medicine because they can be used to control cell fate for multiple therapies, including in regenerative medicine and cancer. However, it remains a major challenge to identify these regulators within the extremely complex and dynamic system of tissues.

Master regulators can be identified using genetic perturbation experiments. High-throughput genetic perturbation screening methods are now available with single-cell gene expression readouts^1,2^. However, none of these studies use cell fate as an output to identify regulators. A number of linear and non-linear computational methods have been developed to predict the effect of specific perturbations on cell fate. Methods have been developed using a range of computational methods to predict gene expression programs that explain the difference between perturbed and unperturbed states^3–5^. Computational methods which determine the ordering of cell states along a trajectory, based on their gene expression profiles using a pseudotime or actual time approach^6–10^, have been used to examine the decision-making process of cells by identifying genes that are differentially expressed between branch points in a cell trajectory. These methods often have troubles identifying accurate trajectories and branch points; furthermore, none of the above methods are designed to specifically identify regulators.

The goal of the present study is to understand how cell-type frequency changes in response to different perturbations, and to identify target genes important in regulating cell fate for each cell type. To accomplish this goal, we develop a computational method, Fatecode, to predict master regulators and key pathways controlling cell fate. Fatecode combines in silico perturbation experiments with cell trajectory modeling using a deep learning-based classification autoencoder and associated energy manifold learning. By examining how cell state switching is affected by systematic perturbations introduced to the model, Fatecode can identify master regulators and key pathways in specific cell trajectories. Unlike other methods which require perturbation data (e.g.,^4,11^), Fatecode uses only scRNA-seq data from wild-type samples to learn how cell type distribution changes after each perturbation, by analyzing the relationship between the learned latent layer, gene space, and cell type space. We assessed Fatecode’s performance using gene expression profiles in blood and developing brain from zebrafish and mouse^12–14^ The method’s ability to detect master regulators is further validated using simulated data produced by a mechanistic model based on the Langevin equation with pre-defined master regulators^15^.

## Results

### Fatecode method overview

Fatecode combines an autoencoder with a classifier. Taking single-cell gene expression profiles as input, the autoencoder learns a latent space with reduced dimensions capturing the input information that is useful for cell type classification. The latent space is also passed to the classifier to identify cell types. Each latent layer node of the autoencoder, representing a reduced dimension of the input, is systematically perturbed to simulate altering key gene expression programs, and cell types are reclassified to characterize the effect. The autoencoder’s decoder generates a gene-by-cell matrix of gene prioritization scores. This matrix can be used to identify genes by their importance for the perturbation effect (see method section and Figure 1) that generates the resulting cell type distribution. This approach is inspired by latent vector operations used in natural language processing and computer vision applications to generate novel text and images^16–19^. In those applications, perturbation operations performed on the latent layer generally yield superior results compared to operations performed directly in the input space. The classification component of Fatecode is used to exclude regions that do not conform to the overall structure of the data. This helps in learning a model which is more representative of the underlying data distribution.

**Figure 1:**
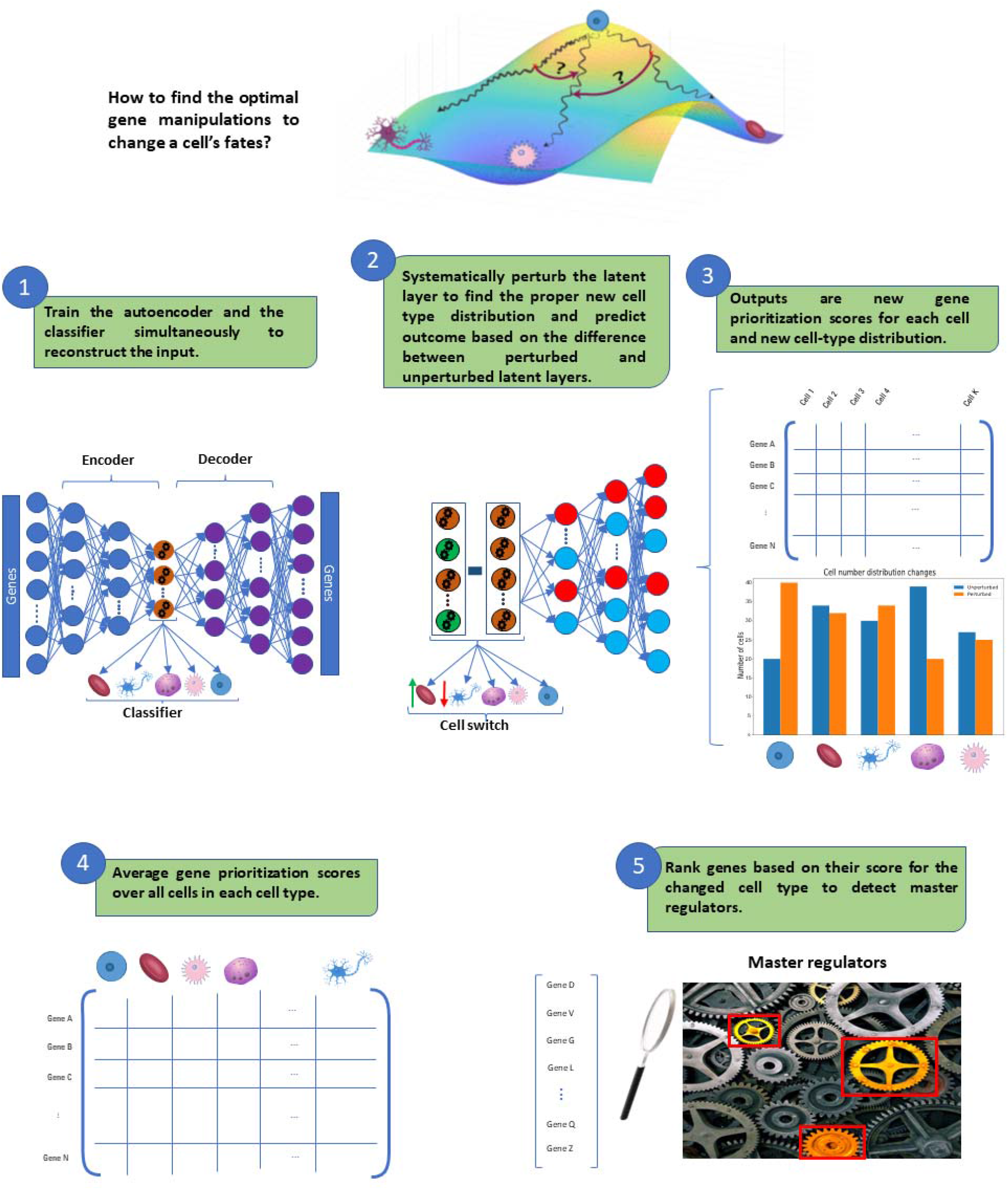
Fatecode workflow for in silico perturbation experiments and master regulator detection. The model is trained with the proper hyperparameter by using gene expression data with a combined cell type classifier and the autoencoder loss function. The latent layer is systematically perturbed and by investigating all possible fate transitions discovered by model perturbation, fate transitions of interest are identified. Perturbation output is simulated by subtracting the perturbed from unperturbed latent layers and feeding it to the decoder part of the autoencoder to identify a cell by gene matrix of prioritization scores that can help us to prioritize genes predicted to be important for cell fate switching. An average of the gene prioritization scores across cells in each cell type is computed. By sorting these genes for our specific cell type, the model predicts genes that should be targeted for cell switching.

### Optimizing model architecture and hyperparameters

Fatecode relies on an autoencoder, but different autoencoder architectures may produce different results, depending on the input data (see supplement)^4, 20–22^. To investigate this in our problem context with single-cell RNA-seq data, we evaluated the performance of three common autoencoder architectures: under-complete autoencoder (AE), variational autoencoder (VAE), and conditional variational autoencoder (CVAE). Thus, the first step of Fatecode evaluates these three available autoencoder architectures to find the one that reconstructs the input data best. Fatecode is implemented in a modular fashion, enabling new autoencoder architectures to be added in the future. To illustrate this feature, we compared how each autoencoder type learns the underlying representation and identifies informative features in the latent layer for single-cell gene expression in adult zebrafish blood^12^. After a hyperparameter search (number and size of all neural network layers and choice of activation function), we chose from each autoencoder class the architecture with the minimum loss (mean squared error, MSE, for reconstruction, and cell type classifier cross-entropy). We then computed the correlation between the input of the encoder and the output of the decoder. AE produced the highest mean (averaged over cell types) correlation and lowest loss (Figs. 2b,d). AE also produces a latent layer that successfully reduces the dimension and cleanly separates the five known cell types in the data (Fig. 2a), and its classifier yields a high accuracy (Fig. 2c).

**Figure 2:**
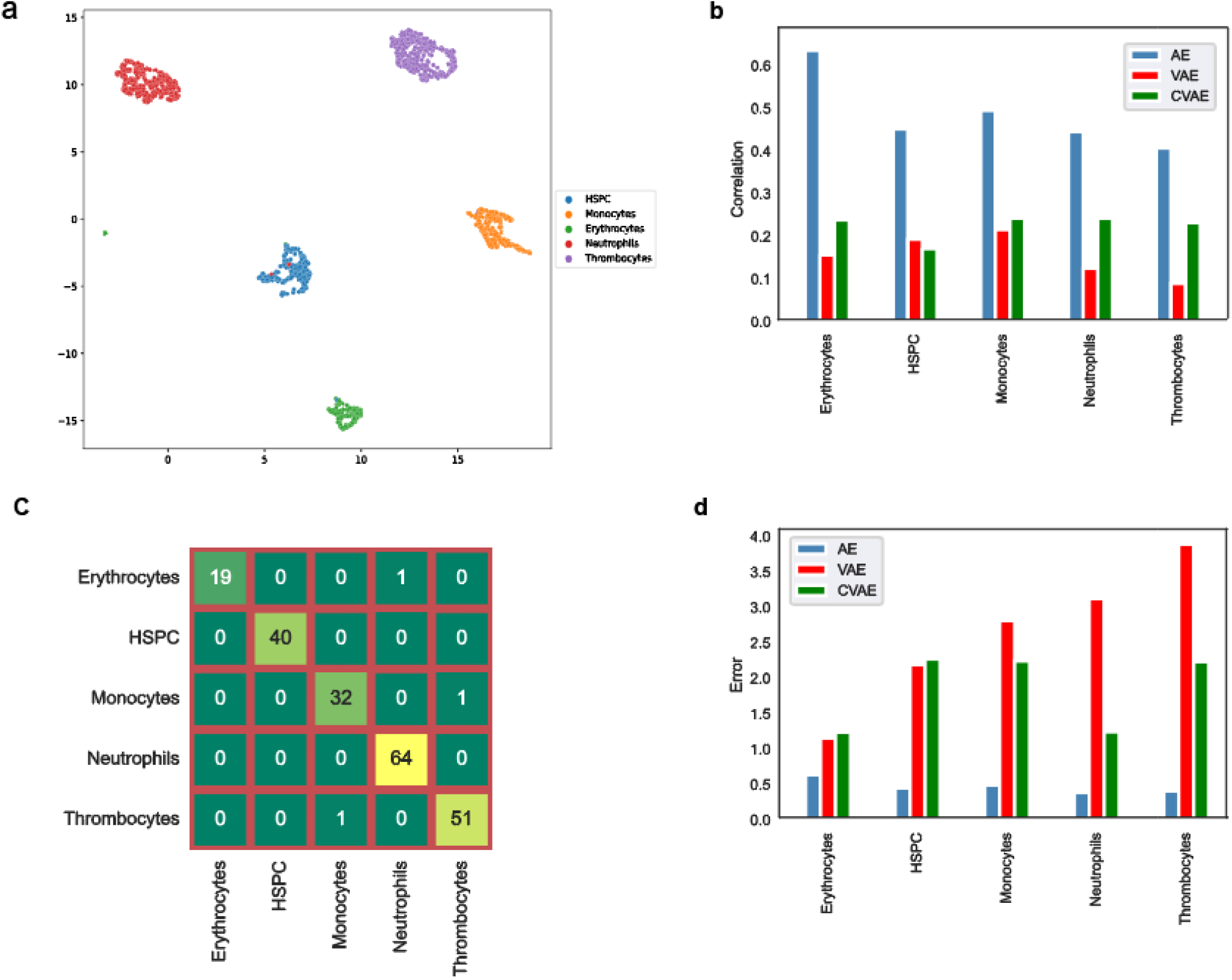
Comparison of classification autoencoder architectures for analyzing data for hematopoiesis regulation in zebrafish. a, UMAP plot visualization of the latent layer of the undercomplete autoencoder (AE). b, comparison of correlation between input and output of AE, variational autoencoder (VAE), and conditional variational autoencoder (CVAE). c, confusion matrix for the classifier connected to the latent layer of AE. d, mean square error between input and output of the three autoencoder architectures. For this dataset, AE has the highest correlation value (b) and lowest error (d), and its latent layer can accurately identify and separate cell types (see supplement).

### Fatecode accurately detects master regulators from simulated scRNA-seq data

To assess the accuracy by which Fatecode identifies master regulators using gene expression profiles, we apply the method to simulated datasets with known gene regulatory network (GRN) structures. We used SERGIO to simulate stochastic gene expression data^15^. SERGIO enables users to specify the number of cell types to be simulated, and the number of master regulators in the GRN (Fig. 3a). A matrix of 400 cells and 2700 genes, with 20 master regulators and 9 cell types, was generated, and run through Fatecode. A set of important genes with their prioritization scores were selected as master regulators, and compared to the known SERGIO list. As expected, the number of known master regulator genes identified increases as more genes are prioritized (Fig. 3b). Remarkably, almost all of the known master regulator genes (18 out of 20) are identified when 150 genes are prioritized (out of 2700).

**Figure 3:**
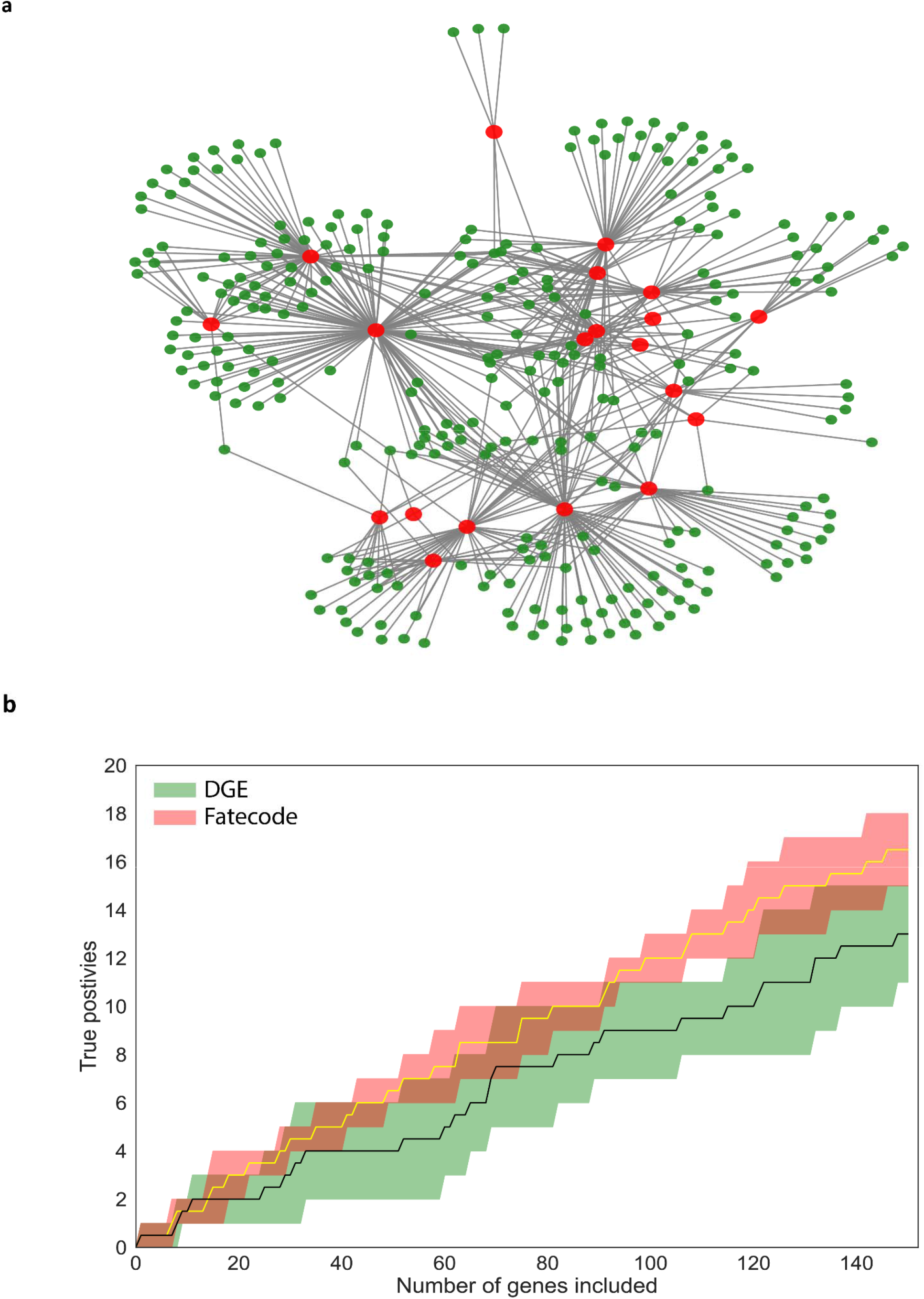
Fatecode detects master regulators using simulated data generated by SERGIO. a, The structure of the gene regulatory network generated by SERGIO. Red nodes are master regulators and green nodes are non-master regulators whose production rates are determined by their associated regulators. b, Benchmark comparisons of the detection rate of predefined master regulators using DGE vs. Fatecode. The red and green areas represent the performance of Fatecode and DGE respectively, on the simulated data.

We then compared our method with the differential gene expression (DGE) analysis, a traditional method for ranking genes from transcriptomics studies. We applied the Seurat package’s default implementation of DGE analysis, which is based on the non-parametric Wilcoxon rank-sum test^23^. 20 differentially expressed genes were selected from each cell type. Fatecode identifies a greater proportion of known master regulators than DGE analysis over up to 150 prioritized genes (Fig. 3b). To demonstrate the robustness of the result, we analyzed five additional datasets generated using SERGIO, all of which yielded similar results (plotted as shading in Fig. 3b). Overall, our simulation analysis indicates the potential of Fatecode for understanding key regulators in GRN, possibly in combination with DGE analysis.

### Fatecode can detect important regulators in cell differentiation and lineage commitment in zebrafish

We applied Fatecode to zebrafish hematopoiesis data^12^. From all possible perturbations on the latent layer performed by Fatecode, we selected ones that resulted in the greatest predicted relative increase in Hematopoietic Stem and Progenitor Cells (HSPCs) (Fig. 4a). As shown in Fig. 4b, following the perturbation, some cells (mostly monocytes) are predicted to switch to HSPCs (Fig. 4b). Fatecode prioritized Signal Transducer And Activator Of Transcription 5A (*stat5a*) as one of the most important genes for this process. It is known as a major regulator of normal hematopoiesis with pleiotropic roles in hematopoietic stem cells^24^ Deletion of *stat5a* leads to an increase in HSPCs cycling, gradually reduced survival, and depleted the HSPCs pool^25^. Another intriguing candidate is *irf8*. It is a key regulator of myelopoiesis in different model organisms and is critical for the development of monocytes and dendritic cells^26,27^. It functions at an early step of the transcriptional program that governs differentiation from myeloid progenitors to monocytes/macrophages and plays a key role in stem cell renewal and maintenance^28 27^ Fatecode also reports a strong connection between *foxo3* and myeloid cell differentiation, consistent with *foxo3* knockout studies, which show a significant increase in granulocyte/monocyte progenitors in the spleen, bone marrow, and blood and enhances short-term hematopoietic stem cell proliferation^29–31^.

**Figure 4:**
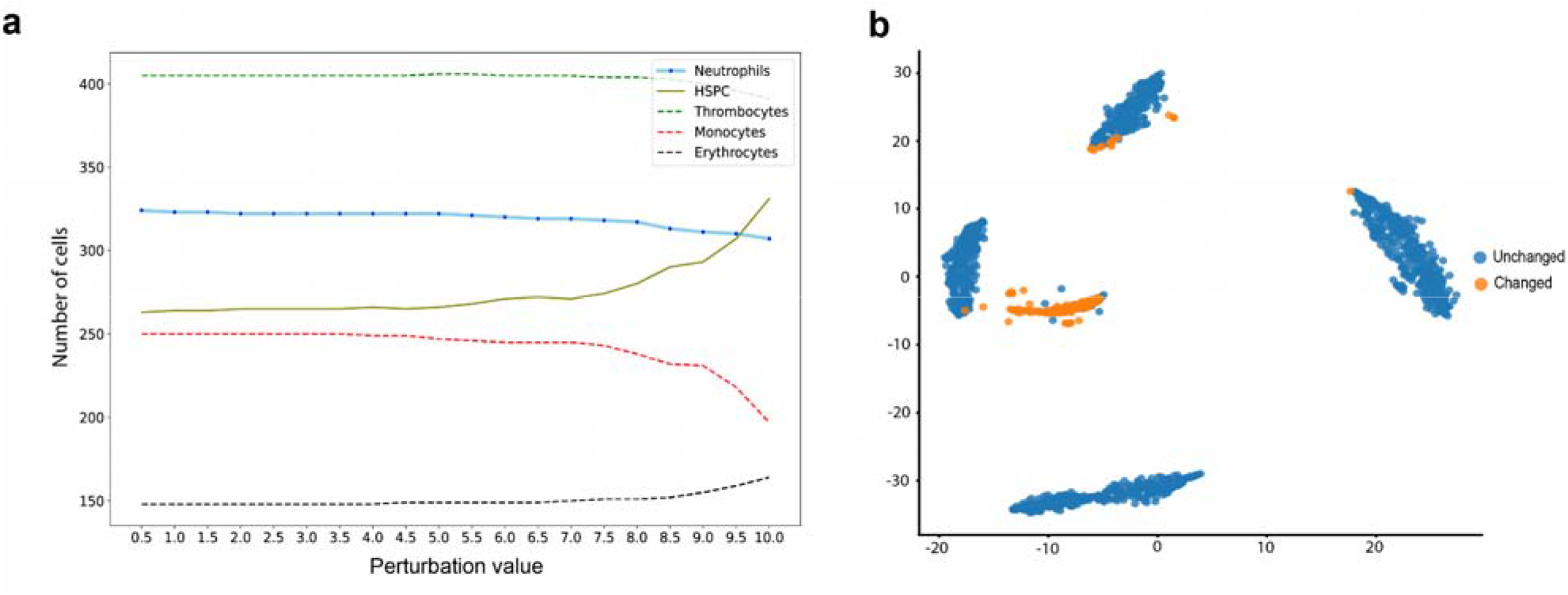
In silico experiments to induce hematopoietic stem/progenitor cells using hematopoiesis in zebrafish. a, A series of latent layer perturbations and their effect on cell distribution. b, Cells that switc**h** from their initial cell type to HSPCs are highlighted.

Fatecode also reveals a relation between *thbs1* and HSPC. *thbs1* is known to limit the expression of key self-renewal transcription factors, including *Oct3* and 4, *sox2, klf4*, and *c-myc* in cells^32^. Other key predicted gene candidates for this perturbation are also known to be involved in hematopoiesis (Table 1).

**Table 1:** List of key gene candidates identified by Fatecode in zebrafish hematopoiesis.

| Gene | Roles | References |
| --- | --- | --- |
| *cdk1* | Plays an important role in the maintenance of pluripotency and genomic stability in human pluripotent stem cells. | 33 |
| *top2a* | Controls the survival of human pluripotent stem cells. | 34 |
| *hmgb2a* | Regulates hematopoietic stem cell maintenance. | 35 |
| *ube2c* | Highly expressed in hESCs and is a biomarker of cancer stemness. | 36,37 |
| *fbxo11* | Its depletion leads to the hematopoietic population with stem cell characteristics | 38 |
| *hmgn2* | Facilitates the maintenance of active chromatin states required for stem cell identity in a pluripotent stem cell model. | 39 |
| *aspm* | Regulates symmetric stem cell division by tuning Cyclin E ubiquitination. | 40 |
| *myb* | Regulates hematopoietic stem cell and myeloid progenitor cell development. | <sup>41</sup> |
| *kpna2* | Exhibits strong interactions with oct4 in embryonic stem cells. | <sup>42</sup> |

### Fatecode reveals key player genes in mouse hematopoiesis

Hematopoiesis is a cell differentiation process by which the body produces blood cells and blood plasma. Here we applied Fatecode to a published mouse hematopoiesis single-cell differentiation dataset that involves differentiation into twelve cell types (Fig. 5a), to examine the method’s accuracy in detecting master regulators in cell differentiation into neutrophils and monocytes^43^. Fatecode revealed one latent node (#10) that, when perturbed, simultaneously increases the neutrophil-monocyte population and inhibits their differentiation into neutrophils and monocytes (see Fig. 5b). Fatecode identified genes with the lowest prioritization scores that are associated with increased neutrophil-monocyte cells and decreased neutrophils and monocytes (Fig. 5c). One such candidate is Entpd8, the deletion of which in mice elevates the neutrophil and monocyte population. Our results are consistent with Tani et al. who reported that *Entpd8^-/-^* mice have a higher number of neutrophils, monocytes, and DCs. Furthermore, hydrolysis of luminal ATP by Entpd8 prevents innate intestinal pathology^44^ Fatecode also reveals Nlrp6 as one of the master regulators in neutrophil and monocyte differentiation. Cai et al. showed that the number of hematopoietic stem cells and granulocyte-monocyte progenitors is reduced in Kp-infected Nlrp6^-/-^ mice, while the survival of matured neutrophils in bone marrow is increased^45^.

**Figure 5:**
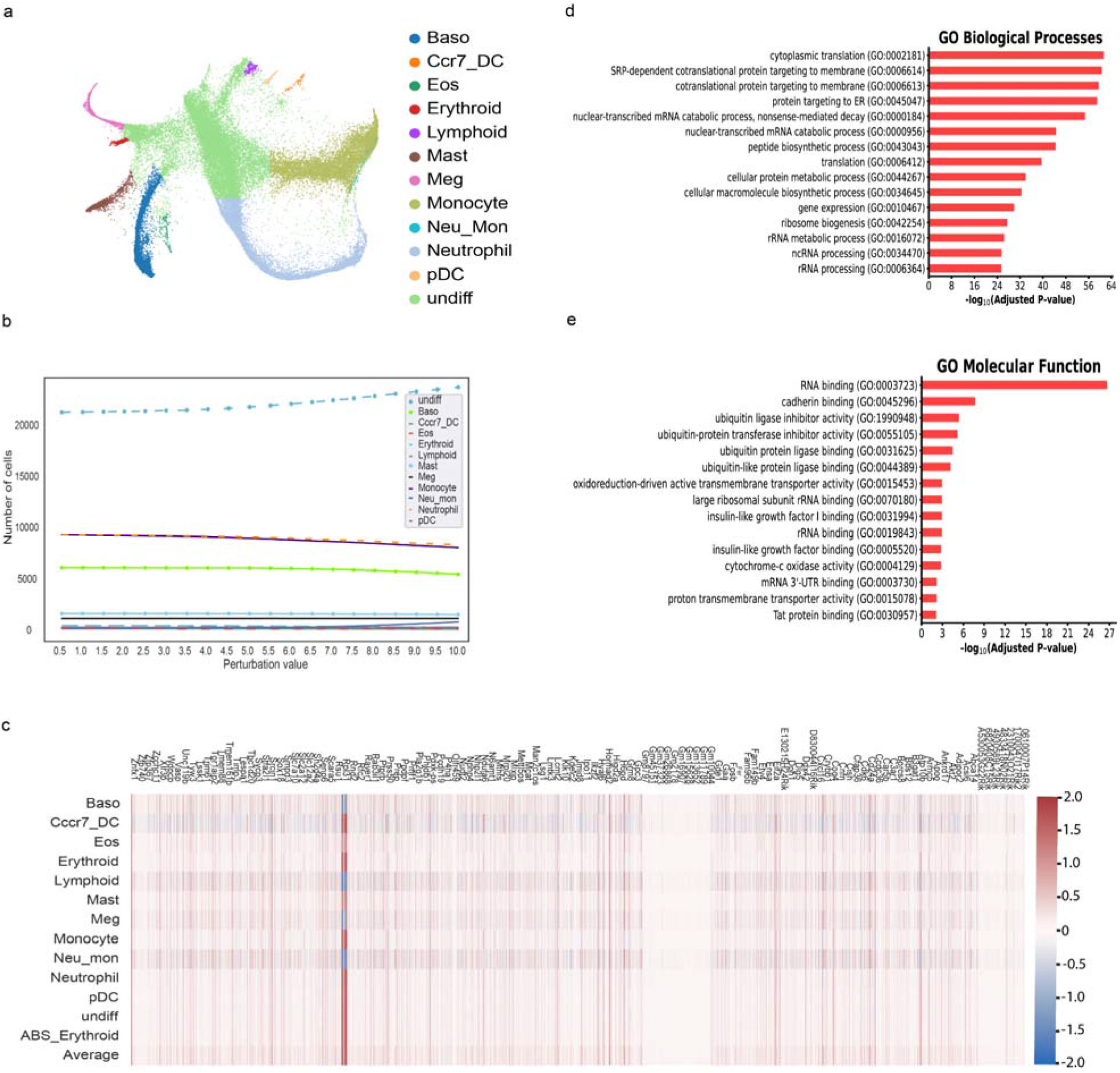
Fatecode accurately detects master regulators and predicts the effect of single-cell perturbation. a, visualization of the hematopoiesis dataset from Weinreb et al.^43^ hematopoietic progenitors differentiate into different cell types such as mast cell (Ma), basophil (Ba), eosinophil (Eos), megakaryocyte (Mk), lymphoid precursor (Ly), migratory dendritic cell (mDC) and plasmacytoid dendritic cell (pDC), erythrocyte (Er), neutrophil (Neu), monocyte (Mo). b, The effect of different perturbation sizes of node #10 on the cell distribution. c, Heatmap of the average prioritization scores of genes for each cell type. d,e, gene set enrichment analysis results. Gene ontology (GO) biological processes and molecular function enrichment analysis show significant process and function terms related to mouse hematopoiesis.

To further understand the role of the top 250 genes in the GRN, gene set enrichment analysis was performed. Pathways that are significantly enriched include neutrophil activation and degranulation, indicating a strong association with the top 250 gene candidates detected by Fatecode (Figs. 5d and 5e).

### Fatecode identifies master regulators in mouse hippocampus development

In the next set of experiments, we applied Fatecode to a dataset for developing mouse hippocampus cells^14^, which is a collection of 18,213 cells and 3,001 genes. The data is clustered in 14 annotated cell types (Fig 6a). We first sought to identify master regulators in the differentiation process that preferentially increases granulocytes (see Fig 6b). Fatecode identifies the ZFP family (Zfp94, Zfp189) and Etv5 as crucial candidates for granulocyte differentiation. Overexpression of Zfp189 in prefrontal cortex neurons is known to preferentially activate the prefrontal cortical, change neuronal activity, and enhance behavioral resilience. The primary upstream regulator in this network is CREB, which binds to Zfp189. It has been previously shown that CREB-Zfp189 interaction in PFC is important for the proliferation, survival, and differentiation of granulocytes (granulopoiesis)^46,47^ The other identified candidate is Etv5. It has been shown that Etv5 is upregulated by HHEX by binding promoter loci, which would be maintained by the ASXL1-MT expression. Expression of either ASXL1-MT or HHEX induces an increase of Etv5, which leads to an increase in granulocyte-monocyte progenitor (GMP)^48^. Other master regulator prioritization scores are shown in Fig. 6c.

**Figure 6:**
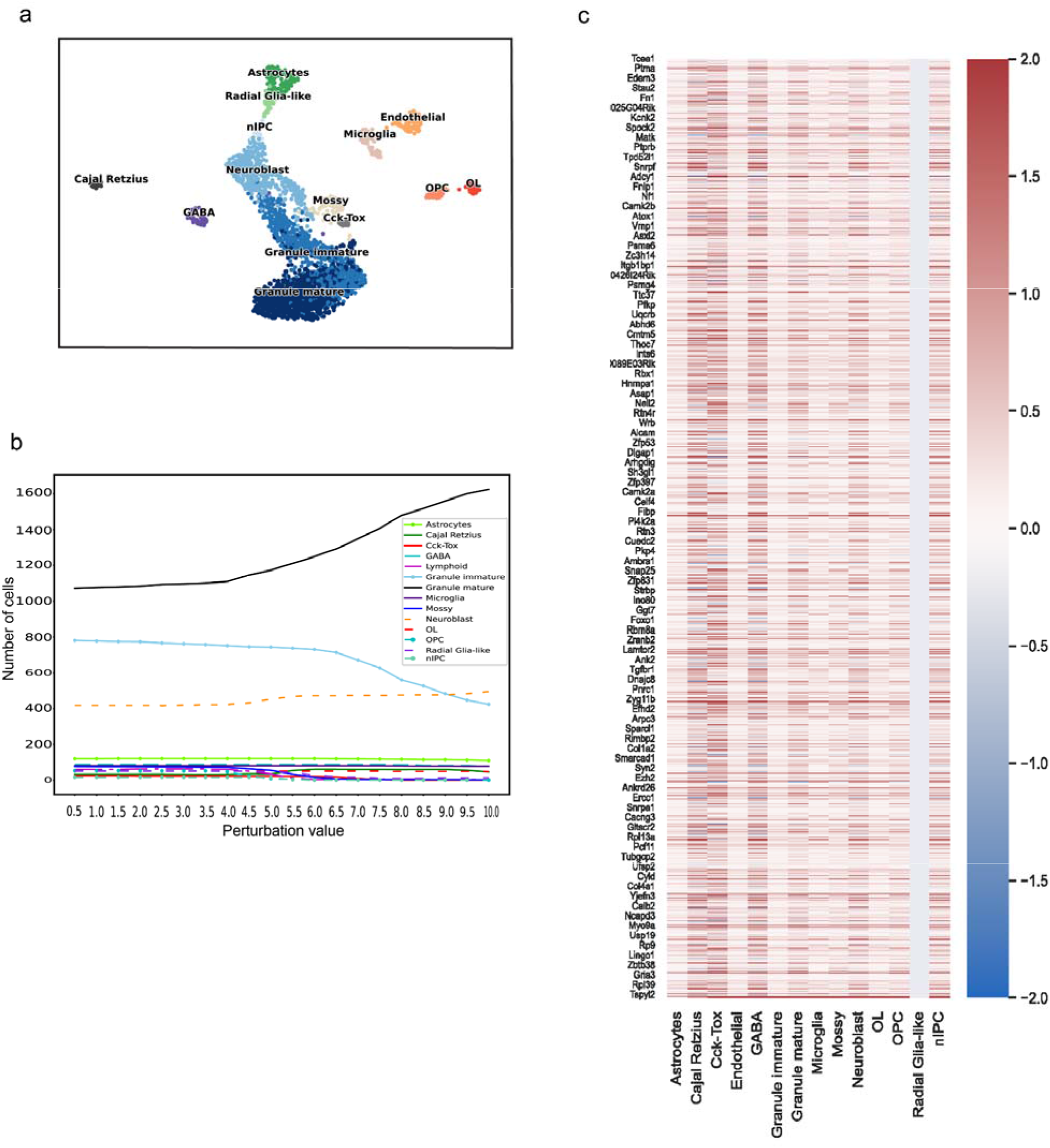
Fatecode identifies key genes in mouse neurogenesis. a, UMAP embedding of fourteen major cell types. b, latent layer node perturbation leads to an increase in granulocytes. c, heatmap of gene prioritization scores for each cell type.

Next, we applied Fatecode to determine master regulators that mediate the differentiation process which preferentially increases oligodendrocyte progenitor cells (OPC), and decreases granulocytes (both mature and immature) and oligodendrocytes. Our model reveals Igfbpl1 as a gene candidate whose expression can have an impact on OPC to oligodendrocytes differentiation which is consistent with RNA velocity analysis and published experimental findings^49,50^. Another candidate is the nuclear receptor 4A subfamily of transcription factors (Nr4a2). Nr4a2 has been linked with inflammation and neuroprotection of the hippocampus, and the expression of Nr4a2 is associated with an increase in quiescent cells. Overexpression of Nr4a2 increases the expression of cell cycle inhibitors such as p18 and p19, both of which are important for G0 to G1 entry^51,52^. Our method correctly predicts that Nr4a2 expression will affect the OPC differentiation into oligodendrocytes and keep the cells in the OPC state. Also, our model selected Smarca, which is part of the SWI/SNF family that is related to chromatin remodelling. Smarca is overexpressed in OPCs and plays an important role during OPC specification and oligodendrocyte differentiation^53,54^.

## Discussion

There has been increasing recognition that the ability to guide the conversion between cell states has tremendous therapeutic value^55^. For such therapies to become a reality, one must identify key genes and optimal paths of these cell state conversions. Therefore, cell reprogramming is a promising technology for tissue repair and regeneration, with the ultimate goal of accelerating recovery from diseases or injuries by manipulating a group of key genes, or master regulators, that together control cell fate. A crucial component in successful cell reprogramming is to correctly identify the master regulators from single-cell transcriptomics datasets. However, the number of genes in these datasets is large and the number of underlying interactions is even larger. Recent studies have demonstrated that a single master regulator’s expression is insufficient to produce an end-point phenotype^56^. Instead, a group of master control networks acts together across a variety of biological processes and pathways to induce a complete lineage conversion^57^ To efficiently and accurately unravel these control networks, we have developed a deep learning method Fatecode, which we have successfully applied to analyze different datasets. First, our method discovers an efficient architecture and latent layer for specific single-cell data. Then by performing operations on the latent layer, it is able to produce useful and potentially novel combinations of perturbations for cell fate reprogramming that could eventually be applied for treatments in disease models and prioritize the predicted gene targets for experimental validation. Genes that have previously been shown in experiments to be crucial in these processes were successfully identified by Fatecode. For further benchmarking we used data generated by the stochastic ODE model and Fatecode was able to detect genes that had been pre-specified as master regulators from the simulated data.

The fundamental idea in Fatecode is similar to the minimum Hamiltonian in physics and potential energy landscape^58^. These authors have shown that the most common autoencoders are naturally associated with an energy function, independent of the training procedure. This reasoning suggests that master regulators can be seen as genes that allow the system to achieve a target celltype redistribution via the most efficient path. Fatecode uses latent layers as a guide to determine what node in the latent layer has to be perturbed. Then the decoder maps the perturbed latent layer to gene space. It’s also useful to understand if master regulators are cell type-specific since some master regulators are crucial for different cell fate decision-making and switching and are not cellspecific. For example, it has been shown that the mammalian target of rapamycin complex (mTORC1) is important in the process of T-cells fate and regulation^59^. Also, mTORC1 regulates the function of most other immune cells, including B cells, neutrophils, monocytes, and macrophages^60–62^. Therefore, in order to achieve a specific phenotype and a successful cell regeneration, subsets of cell-specific and non-specific genes should co-regulate.

Despite offering a useful input data representation, how the autoencoder latent layer represents the input data may be difficult to understand. For example, a gene regulatory program could be represented by a single latent layer node or a set of nodes. The more distributed the information about one gene regulatory program is across nodes, the more challenging it will be to interpret. Future studies will need to better understand how the input data is represented and learned in the latent layer given diverse input data.

In conclusion, we developed an effective computational framework for identifying key players in cell fate control and for predicting the consequences of perturbations on cell distribution. Fatecode’s modular design enables users to select the architecture that would produce an accurate gene detection for their data. By leveraging the power of classification autoencoders and the associated energy manifold learning, Fatecode is a useful tool for answering the critical question of what the most effective combinations of genes are to be manipulated to achieve the desired cell transitions.

## Method

### Deep representation learning

Autoencoders are a class of neural networks with a latent layer capable of learning nonlinear representations of the input data in an unsupervised manner. An autoencoder consists of an encoder that maps the input to the latent space and a decoder which transfers the latent space back to the original space. It can be used for denoising, reducing dimensionality, or learning the representation (or manifold) of the data. We implemented three autoencoder architectures: undercomplete AutoEncoder (AE), Variational AutoEncoder (VAE), and Conditional VAriational Encoder (CVAE) (Fig. 1). AE has a simple latent layer of numerical values, VAE has additional constraints which represent means and standard deviations of multiple Gaussian distributions of the input. By default, the latent layer representation will likely be different each time the representation is learned. The biological task for our autoencoder is to learn a reduced dimension representation of a cell by gene matrix capturing measurements of a single-cell transcriptomics experiment mapping cellular trajectories. Only the gene dimension is reduced, so the latent space describes a reduced representation of each input cell transcriptome. To make the latent layer more specific and reproducible for our biological task, we added a cell-type classification task to the standard regression tasks. The classification task, described in more detail below, predicts the type of each latent cell and compares it to a known input cell type. The training process works to optimize both classification and regression performance. This reduces the space of latent layer candidates since not all possible latent layers are useful for the classification task.

### VAE

VAE is a type of autoencoder that estimates a latent set of probability density functions that model the input data. Unlike AE, which learns an unconstrained representation of the data, VAE assumes that the prior has a Gaussian distribution. An input gene by cell matrix *X* is run through an encoder, which generates parameters for the set of distributions *Q*(*z* | *X*). Then, from *Q*, a latent k-vector *z* is sampled and the decoder transforms *z* into an output, with the condition that the output is similar to the input, where k equals the number of components (or distributions) in the VAE. The VAE total loss consists of the reconstruction loss (first term) and the KL-divergence loss (second term):

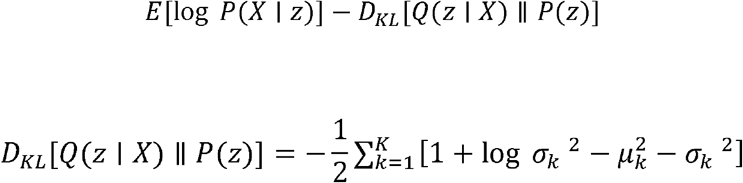

where *μ*_□_ and *σ*_□_ are the k-th components of output vectors *μ_k_*(*X*) and *σ*_□_(*X*), respectively.

### CVAE

CVAE is distinguished from VAE by its embedding of conditional information in the objective function. CVAE relies on two inputs: the features and the class labels, *c*, instead of using only the features, as is done with a VAE and AE. The CVAE architecture allows the encoder and the decoder to be conditioned by *c*. Hence, the variational lower bound objective is changed to the following form:

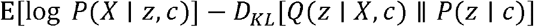

### Overall network architecture of Fatecode

The Fatecode autoencoder architecture was automatically chosen for each of the datasets analyzed in this study using a hyperparameter search (number and size of all neural network layers and choice of activation function). Encoder and decoder architectures are constrained to have the same number of outer and inner layer nodes. For the analysis of hematopoiesis regulation in zebrafish, Fatecode consists of a fully connected encoder and decoder. The encoder and decoder are both two-layer networks of 92 (outer layer) and 48 (inner layer) nodes with the LeakyReLU activation function and the latent layer has 18 nodes. For the analysis of hematopoiesis in mouse data^13^, the encoder/decoder has a 506-node outer layer and a 253-node inner layer, and the latent layer has 125 nodes. For the developing mouse hippocampus data, we used a two-layer encoder/decoder of 50 (outer), 26 (inner), and a latent layer of 15 nodes.

### Classification

The classifier determines cell types using the latent layer as input. The two classifier layers are fully connected. ReLu and softmax activation functions are used for the hidden and output layers, respectively. The number of layers is varied during the hyperparameter optimization; 15 and 5 nodes are chosen for the hidden and output layers, respectively, using adult zebrafish blood data^12^. The classifier has 25 and 12 nodes to classify hematopoiesis in mouse data^13^. For the developing mouse hippocampus data^14^, we select 22 and 14 nodes. It should be mentioned that cell labels are assigned in a self-supervised way using k-means clustering on the latent layer of a separate autoencoder. Also, the clustering labels are compared to the original paper to make sure the cell types are accurate.

### Identifying key regulators in cell differentiation

We have developed a method that analyzes the effects of perturbations on cell fate, by perturbing autoencoder latent variables learned from a single-cell transcriptomics data set capturing cellular trajectories. Consider gene-level adjustments that will morph a baseline cell-type distribution (“A”) to a given target distribution (“B”). To detect genes that are important in this transition, the method proceeds as follows after training:

1. First, the gene expression data, E, associated with all cells with cell-type distribution A are passed through the encoder to generate a matrix of latent variables X. Each column of X is associated with a cell in E; each row corresponds to a latent variable (each variable is represented at a neural network node in the latent layer). X can be used as the input for the classifier to identify each cell’s type.
2. In a series of simulations, finite perturbations of different sizes (e.g., from a 50% reduction to a 10-fold increase) are applied to each row in X sequentially. For each perturbed latent layer X*, the classifier determines a new cell-type distribution.
3. A number of new distributions are generated, one for each latent node for a given perturbation size. Then perturbed layer X* closest to the target distribution B is identified.
4. Next we seek to glean from X* the gene prioritization scores associated with the translation of A to B. To that end, we compute the difference between X* and X. For instance, if increasing latent node #9 k-folds can best approximate B, then the difference between X* and X is a matrix (with rows and columns corresponding to latent nodes and individual cells, respectively) of which the entries that are all zeros, except for the 9^th^ row, which are k times the baseline values.
5. With this “optimal perturbation matrix” (X*-X) computed from the latent layer, the decoder produces a gene-by-cell matrix. Then the average gene expression profile of all cells in each cell type is computed, resulting in a gene by cell_type matrix *M*. The (*i,j*)-th entry of *M* is the prioritization score for the *i*-th gene in cell_type *j*.
6. To identify the master regulators responsible for morphing cell-type distribution A to target B, we consider the cell types whose populations are significantly altered and rank the genes by increasing order of the magnitude of their prioritization scores.

The highly ranked genes (high importance) for a given cell type are those with the lowest prioritization scores. An overall ranking is obtained by calculating for each gene its total score by adding the magnitude of its prioritization score for each relevant cell type and ranking those total scores. It is noteworthy that *M* is the optimal latent node perturbation viewed in gene space and because the perturbation is applied to the latent nodes, *M* does not directly specify how much each gene should be perturbed to yield target B. Nonetheless, *M* contains information that reveals genes that are the most efficient in morphing cell-type distribution from state A to B. This idea is similar to the minimum Hamiltonian and potential energy in physics or the optimal path with the least action^58^.

## Dimensionality reduction for visualization

Python package “UMAP” was used to visualize the latent layer.

## Data preprocessing

The scRNA-Seq gene expression data is log normalized, scaled, and centered.

## Code availability

Codes supporting this study are available on

